# A bifunctional ATPase drives tad pilus extension and retraction

**DOI:** 10.1101/616128

**Authors:** Courtney K. Ellison, Jingbo Kan, Jennifer L. Chlebek, Katherine R. Hummels, Gaёl Panis, Patrick H. Viollier, Nicolas Biais, Ankur B. Dalia, Yves V. Brun

## Abstract

Molecular motors convert chemical energy directly into mechanical work^1^ and are found in all domains of life^2^. These motors are critical to intracellular transport^3^, motility^4,5^, macromolecular protein assembly^3,6^, and many essential processes^7^. A wide-spread class of related bacterial motors drive the dynamic activity of extracellular fibers, such as type IV pili (T4P), that are extended and retracted using so-called secretion motor ATPases. Among these, the tight <u>ad</u>herence (tad) pili are critical for surface sensing, surface attachment, and biofilm formation^8–10^. How tad pili undergo dynamic cycles of extension and retraction^8^ despite lacking a dedicated retraction motor ATPase has remained a mystery. Here we find that a bifunctional pilus motor ATPase, CpaF, drives both activities through ATP hydrolysis. Specifically, we show that mutations within the ATP hydrolysis active site of *Caulobacter crescentus* CpaF result in a correlated reduction in the rates of extension and retraction. Moreover, a decrease in the rate of ATP hydrolysis directly scales with a decrease in the force of retraction and reduced dynamics in these CpaF mutants. This mechanism of motor protein bifunctionality extends to another genus of tad-bearing bacteria. In contrast, the T4aP subclass of pili possess dedicated extension and retraction motor ATPase paralogs. We show that these processes are uncoupled using a slow ATP hydrolysis mutation in the extension ATPase of competence T4aP of *Vibrio cholerae* that decreases the rate of extension but has no effect on the rate of retraction. Thus, a single motor ATPase is able to drive the bidirectional processes of pilus fiber extension and retraction.

## Main text

The best-studied motor ATPases in eukaryotes are required for intracellular transport and motility, and the current dogma is that these motor proteins facilitate transport in a unidirectional manner^1,3^. Consistent with this, individual cells often possess dozens of specialized motor ATPases to facilitate directional movement of specific cargo^3^. Thus, motor ATPase specificity is hypothesized to be important for tight regulatory control over separate operations. In prokaryotes the most widespread motor ATPases are secretion motor ATPases that drive the polymerization of protein subunits into extracellular filaments including type II and type IV secretion systems (T2SS or T4SS), competence pili of Gram-positive bacteria, archaeal flagella, and T4P^6,8,11^. Retraction of the fibers of T4SS^12^ and T4P^8,13–17^ is critical for their function, as is also hypothesized for the T2SS^18^ and Gram-positive competence pili^19^. Despite its importance, the retraction mechanism has remained elusive for most of these systems as they possess a single motor ATPase, which is required for fiber extension (Table 1). Indeed, phylogenetic analysis suggests that the ancestor of these systems possessed a single ATPase^8^. The T4aP represent an exception to this rule, since they possess an antagonistic ATPase, derived through an early duplication of the extension motor ATPase gene^20^, that drives fiber retraction^15,21,22^. How do the majority of these fiber systems retract in the absence of a retraction ATPase?

**Table 1.** Most extracellular filaments that use related extension ATPases to power extension do not possess an antagonistic retraction ATPase.

| Extracellular filament type | Extension ATPase | Predicted retraction/evidence of retraction | Retraction ATPase present? |
| --- | --- | --- | --- |
| Type II secretion | GpsE | Yes | No |
| Gram positive competence pili | ComGA | Yes | No |
| Type IVa pili | PilB | Yes | Yes |
| Type IVb pili | Variable depending on system | Yes | Sometimes |
| Type IVc tad pili | CpaF | Yes | No |
| Type IV secretion | VirB11 | Yes | No |

Here we use the tad type IVc pili (T4cP) as a model to elucidate the retraction mechanism for the systems that lack a retraction ATPase (Fig. S1). Analysis of a local database of fully sequenced bacterial genomes revealed that one third (836/2554) encode an extension ATPase gene that is co-oriented with two platform protein genes, consistent with tad pilus operons^10^ (Fig. 1, Table S2), highlighting that tad ATPases are both highly and broadly distributed even by this conservative metric. We recently showed that the tad pili of *C. crescentus* can retract despite lacking a *cis*-encoded retraction ATPase^8^. We hypothesized that the mechanism of retraction may be mediated by 1) an unidentified retraction ATPase encoded *in trans*, 2) a bifunctional ATPase that powers both extension and retraction, or 3) a process independent of ATP hydrolysis.

**Figure 1.**
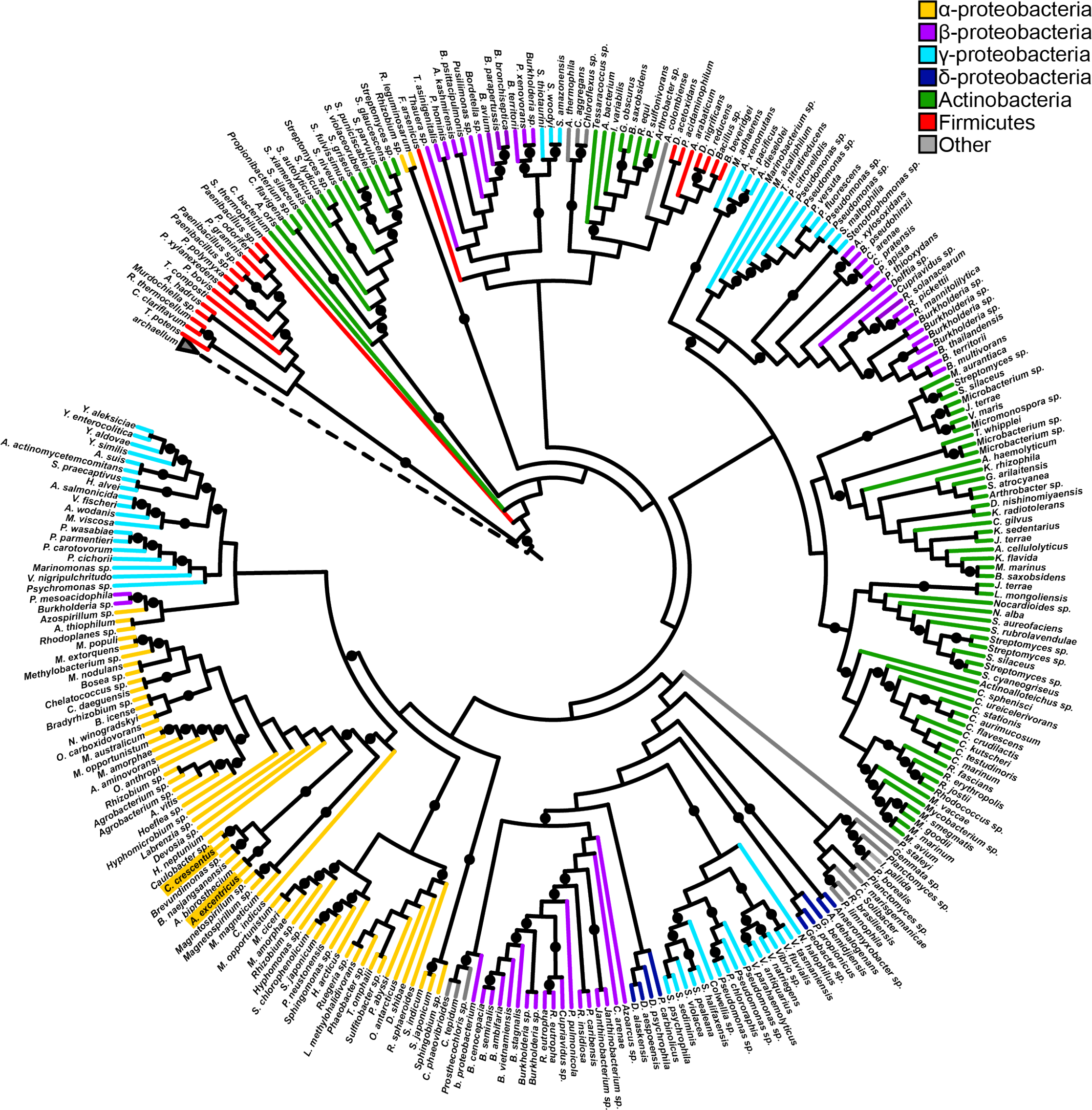
Tad-like ATPases are widely distributed. Rooted cladogram of 297 TadA-like ATPases and 3 archaellum ATPases generated by RAxML maximum likelihood analysis. Branches are colored according to taxonomic class and nodes with rapid boostrap values greater than or equal to 85 are indicated by a black circle. The dashed node represents the archaellum ATPase clade. The accession numbers used to construct the tree can be found in Supplementary Table S2, and the tree can also be viewed at https://itol.embl.de/tree/684565227170291554769353.

To differentiate between these hypotheses, we first sought to identify potential unknown retraction ATPases. Retraction is hypothesized to be critical for bringing pilus-dependent phages in proximity to the bacterial cell envelope where phage entry occurs, and a retraction mutant should hypothetically be resistant to pilus-dependent phage infection. We thus performed transposon sequencing (Tn-seq) on mutant libraries infected with the pilus-dependent phage ɸCb5^23^ to identify genes required for phage infection. A previous study performed similar experiments using the pilus-dependent phage ɸCbK^24^, however we found that pilus retraction was not essential for ɸCbK infection (Fig. S2). In contrast, ɸCb5 infection was prevented when pilus retraction was obstructed (Fig. S2). As expected, Tn-sequencing of infected mutant libraries demonstrated that insertions in the tad pilus-encoding *cpa* genes resulted in increased phage resistance, and furthermore revealed that no additional putative motor ATPase proteins outside of the pilus locus conferred increased phage resistance (Fig. S3, Table S3).

To determine whether the single tad pilus motor CpaF may play a role in retraction, we used a sensitized, hyperpiliated strain of *C. crescentus* that has increased numbers of dynamic pili^25,26^ (Movie S1). Pilus labeling of a ∆*cpaF* complemented mutant confirmed that the expression of the CpaF ATPase is required for pilus extension as shown previously^26^ (Fig 2a). We next assessed the dynamics of CpaF localization during pilus extension and retraction. An mCherry-CpaF fusion (Fig. S4) revealed that the ATPase localized to the base of extending or retracting pili equally (Fig. 2b, c, Movie S2). In some events, we noticed that delocalization of mCherry-CpaF from the base of a retracting pilus fiber correlated with a pronounced reduction in retraction speed (Fig. 2d, Movie S3), suggesting that CpaF may play a role in both pilus extension and retraction.

**Figure 2.**
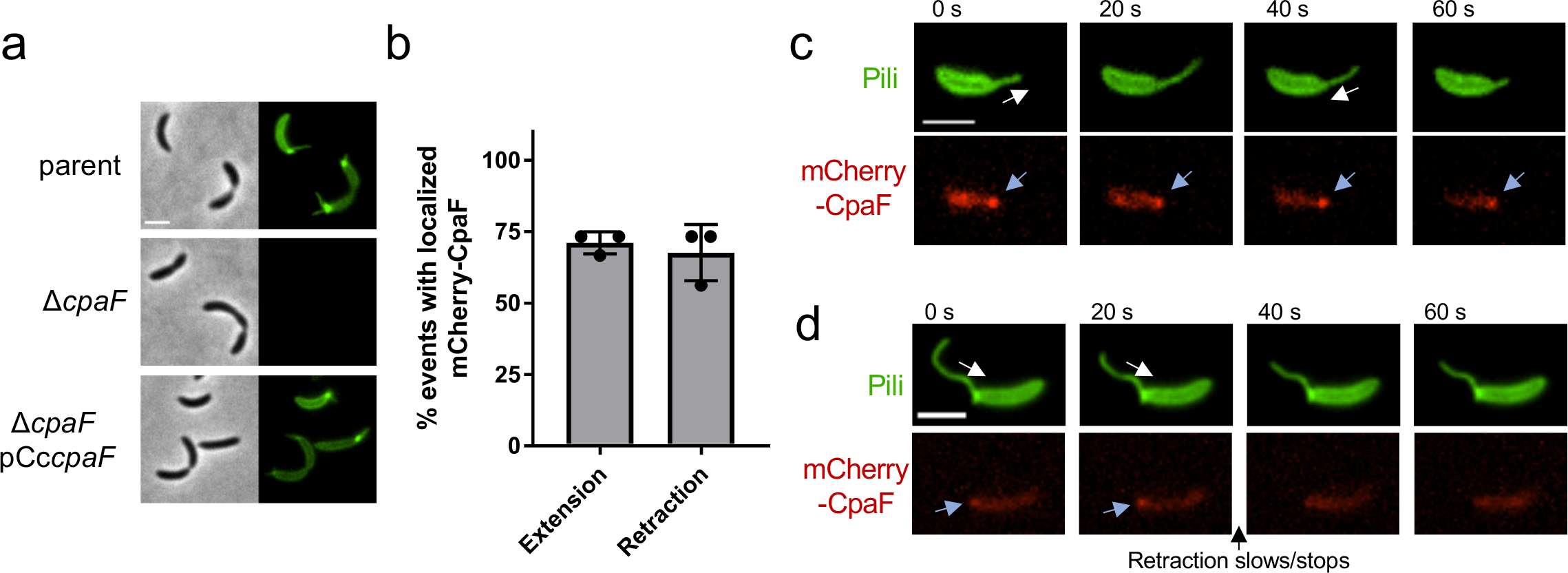
CpaF is required for tad pilus synthesis and is localized during both pilus extension and retraction. (a) Representative images of hyperpiliated CB13 pil-cys strains labeled with AF488-mal. (b) Quantification of pilus extension and retraction events with localized mCherry-CpaF. Data are from 15 extension and retraction events from three independent, biological replicates (n = 45 total extension and retraction events). Error bars show mean + SD. (c) Representative time-lapse images of mCherry-CpaF localization during both pilus extension and retraction. (d) Representative time-lapse images of mCherry-CpaF delocalization during pilus retraction that correlates with halted retraction. Scale bars are 2 µm. White arrows indicate the direction of pilus movement (away from the cell body is extension, towards the cell body is retraction), blue arrows indicate mCherry-CpaF foci.

To distinguish between a bifunctional ATPase model and an ATP-independent model of retraction, we performed an unbiased genetic screen by selecting for retraction-deficient mutants. Because pili are important for adherence^27^ and their retraction is critical for ɸCb5 infection, ultraviolet-mutagenized cells were exposed to ɸCb5 phage and resistant mutants were enriched for those that were still able to attach to surfaces. We then screened mutant isolates for their ability to make pili by fluorescence microscopy. Whole genome sequencing revealed that several isolates contained mutations within the *cpaF* gene (*cpaF*^F244LK245R^ and *cpaF*^D310N^), and Western blot analysis demonstrated that mutant proteins were still expressed (Fig. S1, Fig. S5). Homology-based modeling of CpaF to solved crystal structures^28^ revealed high confidence structural prediction matching a GspE T2SS archaeal ATPase^29^, and mutations identified in CpaF mapped to the predicted ATPase active site of the protein (Fig. S6). These results suggested a role for CpaF ATPase activity in pilus retraction, and we thus made an additional, targeted mutation (*cpaF*^I355C^) that was previously shown to reduce ATPase activity by half in a T4aP retraction ATPase^30^. By fluorescence microscopy, these mutants had reduced extension and retraction rates (Fig. 3a, b, Movies S4, S5, S6), with extension and retraction exhibiting a correlated reduction in rates for each mutation (Fig. S7).

**Figure 3.**
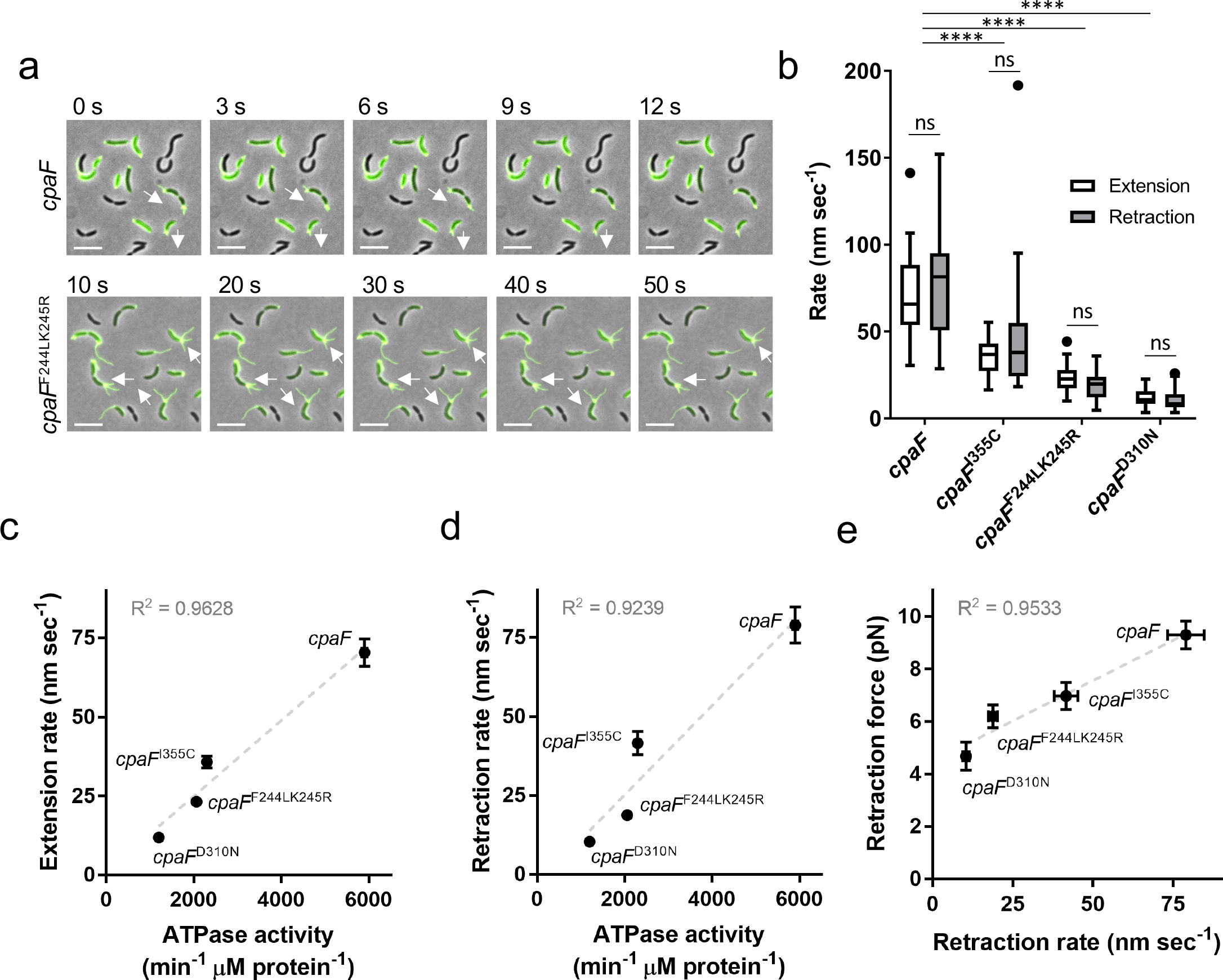
CpaF is a bifunctional ATPase that drives extension and retraction of tad pili. (a) Representative time-lapse images of indicated bNY30a pil-cys *cpaF* strains labeled with AF488-mal. White arrows show directionality of some active pili. Scale bars are 5 µm. (b) Quantification of extension and retraction rates in indicated strains. White boxes show extension rates and gray boxes show retraction rates. Box and whisker plots show 5-95% confidence intervals. Data were collected from three independent, biological replicates. *cpaF* extension n = 30, retraction n = 30; *cpaF*^I355C^ extension n = 30, retraction n = 30; *cpaF*^F244LK245R^ extension n = 30,retraction = 30; *cpaF*^D310N^ extension n = 30, retraction n = 30. Statistics were determined using Sidak’s multiple comparisons test. P****<0.0001, ns = not significant. (c, d) Correlated averages of extension (c) or retraction (d) rates from data shown in (b) and ATPase activity from in vitro ATPase assays. Error bars show SEM. ATPase activity was determined from three replicates of a coupled-enzyme assay where ATPase activity is depicted as the change in NADH per minute per µM protein. (e) Correlated averages of retraction forces and retraction rates. Error bars show SEM. Retraction force measurements of indicated strains were determined from micropillar assays. *cpaF* n = 33, *cpaF*^I355C^ n = 34, *cpaF*^F244LK245R^ n = 34, *cpaF*^D310N^ n = 7.

The above mutations fall into the predicted ATPase active site. We therefore hypothesized that the reduction in extension and retraction rates are a result of altered ATP-hydrolyzing activity. ATP hydrolysis assays of mutant CpaF proteins revealed that they exhibited reduced ATP hydrolysis (Fig. 3c, d). Furthermore, ATP hydrolysis was reduced by varying amounts in different mutants, and this reduction was highly correlated with the reduction in both extension and retraction rates. Together, these data support a model whereby CpaF is a bifunctional motor protein that is driving both extension and retraction through ATP hydrolysis. In line with this, we hypothesized that if CpaF was the motor driving forceful pilus retraction, retraction force should likewise be reduced in ATPase mutants. To measure retraction forces, we employed a micropillar-based assay in which retracting pili bind to elastic micropillars and mediate micropillar bending enabling force measurement^8,31^. *cpaF* point mutants exhibited reduced forces of retraction comparable and correlated to reductions of both ATPase activity and extension/retraction rates for individual mutants (Fig. 3e, Fig. S8), demonstrating that ATP hydrolysis by CpaF drives force generation for tad pilus retraction.

While these data suggest that CpaF drives both pilus extension and retraction in *C. crescentus*, we sought to determine whether this mechanism extends to other tad pili. The tad pili of *Asticcacaulis biprosthecum*, an alphaproteobacterium that harbors bi-lateral stalks, fall into the same phylogenetic clade (Fig. 1) as those of *C. crescentus*. We labeled the tad pili in *A. biprosthecum* and showed that pilus synthesis in this species is also dependent on expression of CpaF (Fig. S9). Unlike *C. crescentus* tad pili, *A. biprosthecum* tad pili were much less dynamic, with the majority of piliated cells harboring static pilus fibers (Fig. 4a, b, Fig. S9, Movie S7). To test whether the differences in pili dynamics were due to differences in CpaF, we performed a cross-complementation experiment where we expressed either the *C. crescentus cpaF* (Cc*cpaF*) in a Δ*cpaF* mutant of *A. biprosthecum* or the *A. biprosthecum cpaF* (Ab*cpaF*) in a Δ*cpaF* mutant of *C. crescentus* (Fig. 4a). Expressing *A. biprosthecum* CpaF in *C. crescentus* reduced the percentage of piliated cells that exhibited dynamic activity by approximately four-fold, while expressing *C. crescentus* CpaF in *A. biprosthecum* resulted in a four-fold increase in the percent of cells harboring dynamic pili, supporting a role for CpaF in the control of tad pilus dynamic activity (Fig. 4b, Movies S8, S9). Because expression of the *C. crescentus* motor ATPase increased the number of cells with active pili in *A. biprosthecum*, we used the Cc*cpaF* and Cc*cpaF*^I355C^ alleles to determine whether decreased ATPase activity would result in reduced extension and retraction rates for this tad system as observed in *C. crescentus*. In line with the model that CpaF mediates both extension and retraction of tad pili, expression of the ATPase “slow” *cpaF*^I355C^ allele resulted in proportional reductions in extension and retraction rates (Fig. 4c). The rate of retraction in *A. biprosthecum* expressing *cpaF*^I355C^ was two-fold higher than in *C. crescentus*, suggesting that other factors play a role in modulating rates of dynamic extension and retraction.

**Figure 4.**
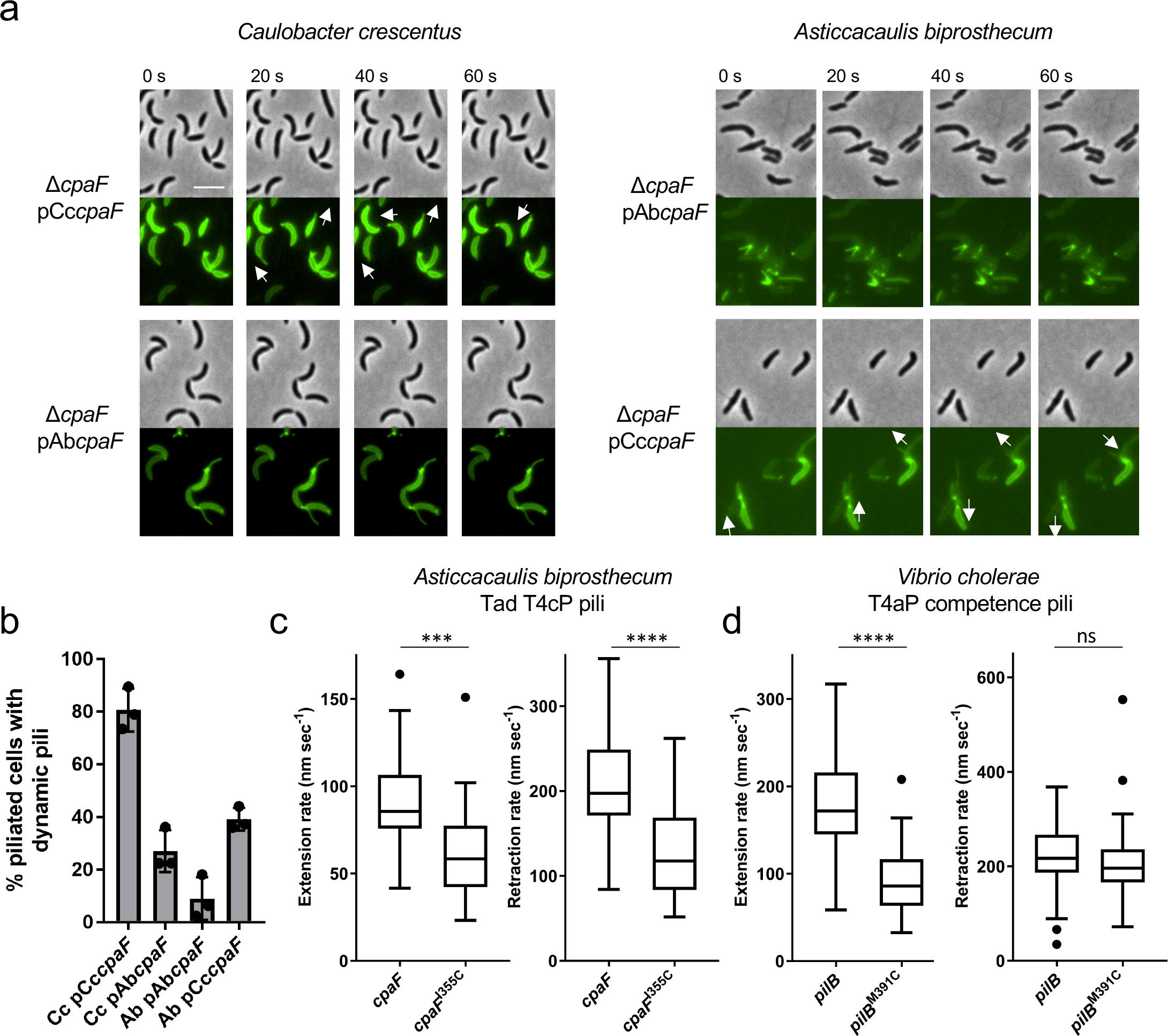
Bifunctional ATPase activity of CpaF is conserved in other tad pilus systems. (a) Representative time-lapse images of indicated pil-cys strains labeled with AF488-mal that contain either their native CpaF motor protein or the cross-complement of the other species (Cc = *Caulobacter crescentus*, Ab = *Asticcacaulis biprosthecum*). White arrows show directionality of some active pili. Scale bar is 4 µm. Look-up tables were normalized for each species to the same values. (b) Quantification of the percent of piliated cells with actively extending and retracting pili for indicated species shown in (a). Number of piliated cells quantified for each strain: Cc pCc*cpaF* n = 163, Cc pAb*cpaF* n = 158, Ab pAb*cpaF* n = 103, Ab pCc*cpaF* n = 161. Bars show mean with SD, and each point represents one replicate. (c) Extension (left panel) and retraction (right panel) rates of *Asticaccaulis biprosthecum* type IVc tad pili using Cc *cpaF* wildtype or I355C mutant alleles. Box and whisker plots show 5-95% confidence intervals. Number of events quantified: *cpaF* extension n = 30, retraction n = 30, *cpaF*^I355C^ extension n = 29, retraction n = 27. (d) Extension (left panel) and retraction (right panel) rates of *Vibrio cholerae* type IVa pili that have either wildtype or a “slow” M391C mutant allele of the PilB extension ATPase. The PilT retraction ATPase in this system was left in tact. Box and whisker plots show Tukey’s confidence intervals. Number of events quantified: *pilB* extension n = 67, retraction = 69, *pilB*^M391C^ extension n = 68, retraction = 71. Statistics were determined using two-tailed t tests. P***<0.001, P****<0.0001, ns = not significant.

The dynamic activity of T4aP is dependent on opposing ATPases that mediate either extension or retraction. We hypothesized that, in contrast to the CpaF mutations described above, “slow” mutations in the active site of the T4aP extension ATPase in these systems should have no effect on retraction rates. We tested this hypothesis using the T4aP of *Vibrio cholerae*, which exhibit dynamic extension and retraction important for natural transformation^17^. These T4aP use an extension ATPase, PilB, or a retraction ATPase, PilT to extend or retract^17,32^. We made a mutation (*pilB*^M391C^) in the ATP-hydrolyzing domain of PilB analogous to the *cpaF*^I355C^ mutation^30^. Consistent with the model that distinct ATPases drive either extension or retraction of T4aP, the “slow” mutation in the extension ATPase reduced the extension rate by half, while the retraction rate remained unchanged (Fig. 4d).

Our findings elucidate the mechanism of retraction in a class of broadly distributed dynamic extracellular filaments and provide insight into the evolution of molecular motors. Retraction of related fibers, which include the Gram-positive competence pili, T2SS, T4SS, and other classes of T4P, is proposed to be essential for their function in fundamental bacterial behaviors including conjugation, natural transformation, motility, secretion, and surface attachment. The majority of these nanomachines possess a single ATPase. Therefore, retraction may be mediated by a bifunctional motor protein in these systems, including the *Vibrio cholerae* toxin co-regulated pili (TCP) where insertion of a minor pilin is thought to initiate retraction^13^. Evolution of a dedicated retraction ATPase in the T4aP and some T4bP may have enabled faster and more forceful retraction. In addition, this evolution may have provided tighter regulation of the extension-retraction switch or provided a deeper regulatory plasticity over pilus retraction frequency, which may otherwise be less flexible and hardwired through a sole biochemical input in single ATPase pilus systems.

## Supporting information

Move S1

## Acknowledgements

We thank S. Shaw, J. Shaevitz, D. Kearns and C. Fuqua for helpful discussions comments on the manuscript. We would like to thank P. Caccamo for the *hfsA* deletion plasmid and advice on *A. biprosthecum* genetics. We also thank R. Snyder for preliminary work on this project. We thank the Center for Genomics and Bioinformatics at Indiana University for whole genome sequencing and SNP mutant identification. This study was supported by grant AI116566 from the National Institutes of Health to NB, by grant R35GM122556 from the National Institutes of Health and by a Canada 150 Research Chair in Bacterial Cell Biology to YVB, by grants R35GM128674 and AI118863 from the National Institutes of Health to ABD, and by National Science Foundation fellowships 1342962 to KRH and CKE.

## Author contributions

CKE designed and coordinated the overall study. CKE designed and performed the CB13 and *A. biprosthecum* experiments. NB designed and JK performed the micropillar experiments. AB designed and JC performed the *V. cholerae* experiments. KRH performed the phylogenetic analysis. PHV and GP designed, and GP performed the Tn-sequencing experiments. All authors analyzed the data. CKE and YVB wrote the manuscript. All authors contributed to editing the manuscript.

## Data availability

The data that support this study are available from the corresponding author upon request.

## Materials and methods

### Bacterial strains, plasmids, and growth conditions

Bacterial strains used in this study are listed in Table S1. A hyperpiliated derivative of *C. crescentus*^25^ was used throughout this study unless otherwise noted. *C. crescentus* strains were grown at 30 °C in peptone yeast extract (PYE) medium^33^ supplemented with 5 µg/ml kanamycin where appropriate. *V. cholerae* strains were grown at 30 °C in lysogeny broth (LB) medium. *Escherichia coli* DH5α (Bioline) was used for cloning and *E. coli* Rosetta2 (Novagen) was used for CpaF overexpression. *E. coli* strains were grown in LB medium at 37 °C supplemented with 25 µg/ml kanamycin and 20 µg/ml chloramphenicol where necessary.

Plasmids were transferred to *C. crescentus* by electroporation or conjugation with S-17 *E. coli* strains as described previously^34^. In-frame deletion strains were constructed by double homologous recombination using pNPTS-derived plasmids as previously described^35^. Briefly, plasmids were introduced to *C. crescentus* and then two-step recombination was performed using kanamycin resistance selection followed by sucrose sensitivity selection. Complementation constructs were constructed using the low-copy number vector pMR10 with genes under control of the leaky *lac* promotor.

For construction of the pNPTS-derived plasmids, ~500 bp flanking regions of DNA on either side of the desired mutations were amplified from bNY30a or *A. biprosthecum* C19 genomic DNA where appropriate. Point mutations were built into the R1 and F2 primers as indicated in Table S1. Upstream regions were amplified using F1 and R1 primers while downstream regions were amplified using F2 and R2 primers. The resulting amplified DNA was purified (Qiaquick) and assembled into pNPTS138 that had been digested with restriction enzyme *Eco*RV (New England Biolabs) using HiFi Assembly Master Mix (New England Biolabs). For the plasmid pNPTS138*cpaF::mCherry-cpaF*, ~500 bp upstream of the *cpaF* gene was amplified from bNY30a genomic DNA using the indicated F1 and R1 primers, the gene encoding mCherry was amplified from vector pRVCHYN-2^36^ using F2 and R2 primers, and the first ~500 bp of the *cpaF* gene were amplified from bNY30a genomic DNA using primers F3 and R3. A codon-optimized DNA sequence encoding a linker (GSAGSAAGSGEF) in frame between the *mCherry* gene and the *cpaF* start codon was built into primers R2 and F3. All three DNA products were then assembled into pNPTS138 as described above.

For construction of pMR10-derived plasmids, primers pMR10CB13cpaFF and pMR10CB13cpaFR were used to amplify the *cpaF* gene from bNY30a genomic DNA and pMR10AbcpaFF and pMR10AbcpaFR were used to amplify the *cpaF* gene from *A. biprosthecum* C19 genomic DNA. For CB13 *cpaF* point mutation alleles *cpaF*^I355C^, *cpaF*^D310N^, and *cpaF*^F244LK245R^, upstream regions of the mutation were amplified using primers pMR10CB13cpaFF + R1 primers and downstream regions of the mutation were amplified using F2 + pMR10Cb13cpaFR primers as indicated in Table S1 with the point mutations built into the R1 and F2 primers. The upstream and downstream DNA were purified (Qiaquick) and then stitched together using splicing by overlap extension (SOE) PCR^37^. *cpaF* regions from CB13 were then digested using restriction enzymes *Sac*I (New England Biolabs) and *Eco*RI (New England Biolabs) followed by heat inactivation at 65 °C for 20 min. The *cpaF* region from *A. biprosthecum* was digested using restriction enzymes *Hind*III (New England Biolabs) and *Eco*RI followed by heat inactivation at 80 °C for 20 min. Digested products were then ligated into plasmid pMR10 that was digested with the same enzymes.

For construction of pET28-derived plasmids, primers pET28CB13cpaFF and pET28CB13cpaFR were used to amplify the *cpaF* gene from bNY30a genomic DNA. For CB13 *cpaF* point mutation alleles, up and downstream regions were amplified as described above with the point mutation built into the R1 and F2 primers and stitched together in the same way using SOE PCR. *cpaF* regions were then digested using restriction enzymes *Nde*I (New England Biolabs) and *Eco*RI followed by heat inactivation at 65 °C for 20 min and ligated into pET28 that was digested with the same enzymes.

*V. cholerae* mutants were constructed by MuGENT and natural transformation as previously described^17,38,39^ and derived from the El Tor isolate E7946^40^. In short, transforming DNA was constructed using splicing-by-overlap extension PCR and transformed or co-transformed into strains^38,39^.

### Phylogenetic analysis

To determine which bacteria encode a Tad-like ATPase, a local database of 2554 completed bacterial genomes were annotated with the Pfam library using the software hmmer v 3.0 and an E value threshold of 1e-10^41,42^. The presence of a TadABC-like operon was established by the presence of 3 co-oriented open reading frames separated by no more than 500 bp that encode proteins containing domains found in the TadABC proteins. Specifically, TadB and TadC were identified by the presence of a single domain with at least 2/3 coverage of the T2SSF domain. TadA homologs were identified by the presence of a T2SSE domain. All genomes analyzed as well as the accession numbers identified as TadABC-like can be found in Supplementary Table S2. Of the 1184 TadABC-like operons that were identified, 297 were randomly selected for phylogenetic analysis and three archaellum ATPases were used as an outgroup (Supplementary Table S2). The TadA and archaellum ATPase sequences were aligned using the default parameters of muscle v 3.8.31. The resulting alignment was used to generate a phylogenetic tree with RAxML version 8.1.3 set to perform 100 rapid bootstraps and subsequent maximum likelihood search using the GAMMA model of rate heterogeneity and JTT substitution model^43^. The resulting tree was visualized using the Interactive Tree of Life visualization software^44^ and can also be viewed at https://itol.embl.de/tree/684565227170291554769353. The root was inferred as the branch separating the archaellum ATPases from the Tad-like ATPases.

### Pilus labeling, microscopy, and analysis

In *C. crescentus* and *A. biprosthecum*, pili were labeled as described previously with some differences^8^. Briefly, cells were grown to an OD_600_ of 0.15-0.3. 100 µl were incubated with 25 µg/ml AlexaFluor488 maleimide (AF488-mal) (ThermoFisher) for 10 min at room temperature. Cells were then pelleted by centrifugation at 5,200 x *g* for one min, washed once with 100 µl PYE, and resuspended in a final volume of 5-10 µl of PYE before one µl of resuspended cells was spotted under a 1% agarose (SeaKem) PYE pad and imaged. Imaging was performed on a Nikon Ti-2 microscope using a Plan Apo 60X objective, GFP and dsRed filter cubes, a Hamamatsu ORCAFlash4.0 camera and Nikon NIS Elements Imaging Software. mCherry-CpaF time-lapses were performed at 10 s/frame, and analysis of localized mCherry-CpaF was performed manually using Nikon NIS Elements Analysis Software. Because *cpaF* mutants exhibited differences in rates of extension and retraction, rates were analyzed from strain YB9040 imaged at 3 s/frame, YB9202 imaged at 5 s/frame, YB9241 imaged at 10 s/frame, and YB9240 imaged at 20 s/frame to avoid phototoxicity and photobleaching while still capturing extension and retraction events. Rates were analyzed from strain YB9233 imaged at 3 s/frame and YB9258 imaged at 5 s/frame. Only extension and retraction events that lasted for multiple frames were analyzed. To calculate rates of retraction, the change in pilus length was manually measured using Nikon NIS Elements Analysis Software and divided by the amount of time over which the length change occurred.

In *V. cholerae*, pili were labeled as described previously with some differences^17^. To constitutively activate competence, the master competence regulator TfoX was ectopically overexpressed (via P_*tac*_ -*tfoX*) and quorum sensing was constitutively activated via deletion of *luxO*^38,45^. Strains were grown to the late-log phase in LB Miller broth + 100 µM IPTG (to induce TfoX expression) + 20 mM MgCl_2_ + 10 mM CaCl_2_. Approximately 10^8^ CFUs of this culture were centrifuged at 18,000 x *g* for one min and then resuspended in instant ocean medium (7 g l^−1^; Aquarium Systems) before labeling with 25 μg/ml AF488-mal for 15 min in the dark. Labeled cells were centrifuged, washed twice and resuspended using instant ocean medium. All imaging was performed under 0.2% Gelzan (Sigma) pads made with instant ocean medium. A Nikon Ti-2 microscope with a Plan Apo 60X objective, a GFP filter cube, a Hamamatsu ORCAFlash4.0 camera and Nikon NIS Elements Imaging Software was used to image cell bodies and fluorescently labeled pili. To determine the rates of extension and retraction, labeled cells were imaged by time-lapse microscopy every second for one min. Extension and retraction events were manually calculated using measurement tools of the NIS Elements analysis software. For extension and retraction rate calculations, only cells that began and completed extension or retraction events within a one-min window were analyzed. Pili that were already extending or retracting when imaging began and/or pili that were shorter than 0.3 μm were not analyzed.

### Genome-wide transposon mutagenesis coupled to deep sequencing (Tn-Seq)

Transposon mutagenesis of *C. crescentus* NA1000 was done by intergeneric conjugation from *E. coli* S17-1 *λ*pir harbouring the *himar1*-derivative p*MAR2xT7*^46^. A Tn-library of >200,000 gentamicin- and nalidixic-acid-resistant clones was collected. NA1000::Tn(Gent^R^) bank was grown overnight in PYE and then freshly restarted in PYE either in the absence or presence of bacteriophage φCb5 (at 10^3 MOI). After 8h or 24h of incubation under agitation at 30°C, each cell cultures were harvested, and chromosomal DNA was extracted. Genomic DNA was used to generate barcoded Tn-Seq libraries and submitted to Illumina HiSeq 4000 sequencing (Fasteris SA). Tn insertion-specific reads (150 bp long) were sequenced using the himar-Tnseq2 primer (5’-AGACCGGGGACTTATCAGCCAACCTGT-3’). Specific reads attesting an integration of the transposon on a 5’-TA-3’ specific DNA locus were sorted (Rstudio_V1.1.442) from the tens of million reads generated by sequencing, and then mapped (Map_with_Bowtie_for_Illumina_V1.1.2) to the *C. crescentus* NA1000 genome (NC_011916.1) using the web-based analysis platform Galaxy (http://usegalaxy.org). Using Samtool_V0.1.18, BED file format encompassing the Tn insertion coordinates were generated and then imported into SeqMonk V1.40.0 (www.bioinformatics.babraham.ac.uk/projects/) to assess the total number of Tn insertion per chromosome position (Tn-insertion per millions of reads count) or per coding sequence (CDS). For CDS Tn-insertion ratio calculation, SeqMonk datasets were exported into Microsoft Excel files (Dataset S1) for further analyses, as described previously^47^. Briefly, to circumvent ratio issues for a CDS Tn-insertion value of 0 and CDS that do not share sufficient statistical Tn-insertions, an average value of all CDS-Tn insertions normalized to the gene size was calculated, and 1% of this normalized value was used to correct each CDS-Tn insertion value.

### Identification of mutants deficient in pilus retraction

To isolate mutants deficient in retraction, stationary phase cultures of the parent strain YB9034 were first mutagenized using ultraviolet (UV) light irradiation resulting in at least 90% killing. Approximately 1 × 10^8^ CFUs were then mixed with ɸCb5 phage at a multiplicity of infection (MOI) of one and added to three ml PYE in a 12-well polystyrene plate and incubated at room temperature overnight shaking at 150 rpm on an orbital shaker. The next morning, the medium was removed from the plates, and the wells were washed once with 3 ml of water to remove unattached cells. Three ml PYE were added to each well, and plates were then incubated again shaking at room temperature for 3.5 hours before an they were washed again. Plates were then incubated shaking for three days until turbidity was observed in the wells. The contents of each well were then struck out onto PYE plates to isolate for single colonies. Isolates were then imaged by microscopy for altered pilus phenotypes and selected for whole genome sequencing.

### Whole genome sequencing and analysis of isolated mutants

DNA was extracted from mutant isolates using the Wizard Genomic DNA Purification Kit (Promega). Briefly, 1.5 ml of stationary phase cultures were centrifuged at 16,000 x *g* for three min to pellet cells. The supernatant was discarded, and pellets were resuspended in 200 µl of TES buffer (10 mM Tris-HCl pH 8, 20 mM EDTA, 1% w/v sarkosyl). RNAse was added to the cell suspension at a final concentration of 0.05 mg/ml and incubated at 70 °C for 10 min. 20 µg of proteinase K was then added to the cell suspension and incubated 70 °C for 25 min. 30 µl of 7.5 M ammonium acetate was then added to the cells. One ml of resin provided in the kit was added to the cell suspension and then flushed through the Wizard Minicolumns. Two ml of Column Wash Solution was then flushed through the columns to clean the DNA. Cleaned DNA was eluted using 80 °C TE buffer (10 mM Tris-HCl pH 8, 1 mM EDTA) and sent to the Center for Genomics and Bioinformatics core facility, Indiana University Bloomington for library preparation and next generation sequencing analysis.

500 ng DNA was sheared on Covaris E220 sonicator and used in library preparation using NEXTflex Rapid DNA-Seq kit (Bio Scientific) according to manufacturer’s protocol. 2x 8 nt dual indexed adapters from TruSeq RNA CD index kit (Illumina) were added to the libraries for multiplexing. The barcoded libraries were cleaned by double side bead cut with AMPure XP beads (Beckman Coulter), verified using Qubit3 fluorometer (ThermoFisher) and 2200 TapeStation bioanalyzer (Agilent Technologies), then pooled. The pool was sequenced on NextSeq 500 (Illumina) with NextSeq75 High Output v2 kit (Illumina) and paired end 2x 38 bp reads were generated. The reads were de-multiplexed using bcl2fastq (software v2.20.0.422) and about 4-6M reads were assigned to each library. Trimmomatic^48^ (version 0.33; non-default parameters) was used to trim reads of adapter and low-quality bases. Reads were mapped to the *Caulobacter vibrioides* strain CB13B1a (CP023315.3) with breseq (version 0.30.1) using default parameters^49^. Custom python scripts were used to combine the breseq output and identify common variants across isolates for a given reference genome. Reads were also mapped to the reference genome with bowtie2^50^ and visualized with JBrowse^51^ to look for complex rearrangements.

### Phage sensitivity assays

Phage sensitivity assays on plates were performed by spotting dilutions of ɸCb5 phage onto lawns of growing *C. crescentus* strains. To make lawns, 200 µl of stationary phase cultures were mixed with three ml of top agar (0.5% agar in PYE) and spread over 1.5% PYE agar plates. After the top agar solidified, five µl of phage diluted in PYE was spotted on top. Plates were grown for two days at 30 °C before imaging.

Phage sensitivity assays in tubes were performed by adding ~10^6^ cfu of indicated strains with ~10^11^ pfu of ɸCb5 phage for a MOI of 10^5^ in three ml PYE where indicated. A final concentration of 500 µM methoxypolyethylene glycol maleimide, MW 5000 (PEG5000-mal) (Sigma) was added to tubes where indicated to block pilus retraction as shown previously^8,52^. Cultures were then grown shaking at 30 °C overnight before imaging.

### CpaF antibody production

For antibody production, an N-terminally tagged full length version of CpaF from *C. crescentus* NA1000 was expressed and purified. Antibodies were raised in New Zealand white rabbits.

### Western analysis

To compare expression of *cpaF* alleles between strains, ~10^9^ cells from exponential phase cultures were centrifuged for three min at 16,000 x *g*. The supernatant was discarded, and pellets were resuspended in PBS (phosphate buffered saline) and SDS loading buffer (62.5 mM Tris-HCl pH 6.8, 10% v/v glycerol, 2% w/v SDS, 0.05% v/v β-mercaptoethanol, 0.0025% w/v Bromophenol blue) was added. Samples were heated to 100 °C for six min and then separated on 10% SDS-polyacrylamide gels. Samples were transferred to nitrocellulose membranes and probed with affinity purified α-CpaF antibody at 1:500 dilution in 5% w/v non-fat milk powder resuspended in 1x TTBS (20 mM Tris-HCl pH 7.6, 130 mM NaCl, 0.05% v/v Tween20) overnight. The nitrocellulose membrane was washed three times with 1x TTBS and then probed at a 1:20,000 dilution of HRP-conjugated goat α-rabbit antibody (Biorad) in 5% non-fat milk solution for one hour. Membranes were developed with Supersignal West Dura substrate (ThermoFisher).

### Force measurement using micro pillars

Retraction forces were obtained by Polyacrylamide MicroPillars assays as described previously^31^. Briefly, an equidistant (3 μm from center to center) polyacrylamide micropillars array with a spring constant of 25+/− 4 pN/μm was obtained by unmolding a silica mold at the center of a 25 mm diameter round coverglass (Warner Instruments). To enable attachment of the pili to the pillars, the pillars were coated with a 0.01% poly-L-Lysine solution (SIGMA) covalently linked to the pillars by UV activated SulfoSANPAH (ThermoFisher) for one hour followed by a one-hour incubation with a water suspension of 0.2% w/v 0.02 *μ*m carboxylate-modified beads (Molecular Probes). After 2 hours of treatment, the cover glass was set in the bottom center of an observation chamber (Invitrogen), then 500 μl fresh PYE liquid medium supplemented with 5 µg/ml kanamycin was added into the chamber. 100 μl mid-log phase bacteria culture was centrifuged at 7,500 rpm for 1 min, the supernatant was removed, and the pellet was resuspended in 100 μl fresh PYE medium containing kanamycin. Then 25 μl of this cell suspension was added to the micropillars array. Subsequently, 10 Hz videos of the pillar tips movement were recorded with a 60x objective. Finally, a combination of ImageJ plugin and Matlab program (Mathworks Inc. Natick, MA) implementing a cross correlation tracking of the pillars’ tip motion was used to analyze the videos and extract the maxima of the pillars’ deflections. Retraction forces were calculated based on pillar calibration.

### Expression and purification of His_6_-CB13CpaF and ATPase assays

250 ml flasks containing 50 ml cultures of Rosetta2 expression strains carrying pET28 plasmids were grown shaking at 37 °C to exponential growth phase OD_600_ (0.5-0.7). Cultures were cooled down on ice for 20 min before 0.1 mM IPTG was added to induce protein expression. Induced cultures were incubated shaking at 16 °C overnight. The following day, 30 ml of cultures were centrifuged at 5,000 x *g* for 10 min at 4 °C to harvest cells. The supernatant was removed, and cells were resuspended in 10 ml Buffer A (20 mM Tris-HCl pH 8, 150 mM sodium citrate, 5 mM 2-mercaptoethanol) with a protease inhibitor tablet (Pierce, EDTA-free). Cell suspensions were sonicated to lyse cells and then centrifuged at 100,000 x *g* to separate soluble and insoluble fractions. Cell lysates were then incubated at 4 °C for 1-1.5 hours on one ml of Ni-NTA resin (Qiagen). The resin was then loaded into columns (Biorad) and washed with 10 ml cold Buffer A. Proteins were eluted with five ml of Elution Buffer (20 mM Tris-HCl pH 8, 150 mM sodium citrate, 5 mM 2-mercaptoethanol, 500 mM imidazole) and collected in 0.5 ml fractions. 2.5 ml of fractions containing the highest protein content were pooled and the imidazole was removed by running the 2.5 ml protein suspension on PD-10 size-exclusion columns (GE Healthcare) followed by cold 3.5 ml of Buffer A to elute proteins. Visual inspection of SDS-PAGE gels indicated that the proteins were >95% pure.

ATP hydrolysis activity was measured using a coupled-enzyme assay^53^. 60 µl of ATPase buffer (50 mM Tris-HCl pH 8, 1 mM dithiothreitol, 90 mM NaCl, 10 mM MgOAc, 50 µg/ml bovine serum albumin, 5% glycerol) containing 10 µg protein was mixed with 60 µl of ATPase buffer supplemented with 10 mM ATP, 10 mM MgCl_2_, 1 mM phosphoenolpyruvate, 0.8 mM NADH, 0.6 units of pyruvate kinase, and 0.96 units of lactate dehydrogenase in a 96-well polystyrene plate and incubated at 30 °C. Reactions were performed in triplicate, and absorbance at 340 nm was measured every minute for 1.5 hours. The slope of linear absorbance decay between 25 to 45 min was used to calculate ATP hydrolysis activity for all samples. An NADH absorbance standard curve was used to calculate concentrations of NADH, and ATPase activity rates are reported as the change in nM of NADH min^−1^ µM protein^−1^. Background rates of NADH oxidation were subtracted by normalization of absorbance at 340 nm over time to no protein controls.

### Statistics

Statistical significance was calculated using tests on Prism 7 software. Statistical differences between two groups were calculated using two-tailed Student’s t-tests. Statistical tests between multiple groups were calculated using Sidak’s multiple comparison’s test. Sample sizes were chosen based on historical data and no methods were used to predetermine sample size.

