## Supplemental material for "A bifunctional ATPase drives tad pilus extension and retraction"

**Table S1.** Strains, plasmids, and primers used in this study.

| Strain | Description or construction | Source or reference |
|---|---|---|
| ***E. coli* strains** | | |
| YB9012 | S17-1 / pNPTSCB13*pilA2*$^{T36C}$ | This study |
| YB9026 | S17-1 / pNPTSCB13Δ*hfsDAB* | This study |
| YB8483 | α Select / pNPTS138CB13*cpaF*::*mCherry-cpaF* | This study |
| YB8475 | S17-1 / pNPTS138CB13Δ*cpaF* | This study |
| YB8487 | S17-1 / pMR10Cc*cpaF* | This study |
| YB9200 | α Select / pMR10Cc*cpaF*$^{I355C}$ | This study |
| YB9234 | α Select / pMR10Cc*cpaF*$^{D310N}$ | This study |
| YB9236 | α Select / pMR10Cc*cpaF*$^{F244LK245R}$ | This study |
| YB9051 | α Select / pMR10Ab*cpaF* | This study |
| YB8471 | α Select / pNPTSAb*pilA*$^{G36C}$ | This study |
| YB9201 | α Select / pNPTS138AbΔ*cpaF* | This study |
| YB442 | α Select / pET28CB13*cpaF* | This study |
| YB9257 | α Select / pET28CB13*cpaF*$^{I355C}$ | This study |
| YB9042 | α Select / pET28CB13*cpaF*$^{F244LK245R}$ | This study |
| YB9041 | α Select / pET28CB13*cpaF*$^{D310N}$ | This study |
| Rosetta2 expression strain | $F^-$ *ompT hsdS*$_B$($r_B^-$ $m_B^-$) *gal dcm* (DE3) / pLysSRARE2 ($Cm^R$) | Novagen |
| ***Caulobacter* and *Asticcacaulis* strains** | | |
| NA1000 | *C. crescentus* lab adapted strain | [54] |
| bNY30a | Hyperpiliated derivative of CB13B1a, originally SW16-Pil 200 | J. Poindexter, [25] |
| YB9017 | bNY30a *pilA2*$^{T36C}$ (conjugated strain YB9012 with strain bNY30a) | This study |
| YB9034 (parent) | bNY30a Δ*hfsDAB pilA2*$^{T36C}$ (conjugated strain YB9026 with YB9017) | This study |
| YB8482 (Δ*cpaF*) | bNY30a Δ*hfsDAB pilA2*$^{T36C}$ Δ*cpaF* (conjugated strain YB8745 with strain YB9034) | This study |
| YB9050 | bNY30a Δ*hfsDAB pilA2*$^{T36C}$ *cpaF*::*mCherry-cpaF* (electroporated plasmid from strain YB8483 into strain YB9034) | This study |

| | | |
|---|---|---|
| YB9249 | bNY30a Δ*hfsDAB pilA2*$^{T36C}$ pMR10 (electroporated plasmid pMR10 into strain YB9034) | This study |
| YB9250 | bNY30a Δ*hfsDAB pilA2*$^{T36C}$ Δ*cpaF* pMR10 (electroporated plasmid pMR10 into strain YB8482) | This study |
| YB9040 (*cpaF*) | bNY30a Δ*hfsDAB pilA2*$^{T36C}$ Δ*cpaF* pMR10Cc*cpaF* (mated strain YB8487 with strain YB8482) | This study |
| YB9202 (*cpaF*$^{I355C}$) | bNY30a Δ*hfsDAB pilA2*$^{T36C}$ Δ*cpaF* pMR10Cc*cpaF*$^{I355C}$ (electroporated plasmid from strain YB9200 into strain YB8482) | This study |
| YB9240 (*cpaF*$^{D310N}$) | bNY30a Δ*hfsDAB pilA2*$^{T36C}$ Δ*cpaF* pMR10Cc*cpaF*$^{D310N}$ (electroporated plasmid from strain YB9234 into strain YB8482) | This study |
| YB9241 (*cpaF*$^{F244LK245R}$) | bNY30a Δ*hfsDAB pilA2*$^{T36C}$ Δ*cpaF* pMR10Cc*cpaF*$^{F244LK245R}$ (electroporated plasmid from strain YB9236 into strain YB8482) | This study |
| YB9194 (Δ*cpaF* pAb*cpaF*) | bNY30a Δ*hfsDAB pilA2*$^{T36C}$ Δ*cpaF* pMR10Ab*cpaF* (electroporated plasmid from strain YB9051 into strain YB8482) | This study |
| YB642 | *Asticcacaulis biprosthecum* C19 | [55] |
| YB425 | *Asticcacaulis biprosthecum* C19 *pilA*$^{G36C}$ (electroporated plasmid from strain YB8471 into strain YB642) | This study |
| YB434 (Ab parent) | *Asticcacaulis biprosthecum* C19 Δ*hfsA pilA*$^{G36C}$ (electroporated plasmid pNPTS138AbΔ*hfsA* into strain YB425) | This study |
| YB9209 (Ab Δ*cpaF*) | *Asticcacaulis biprosthecum* C19 Δ*hfsA pilA*$^{G36C}$ Δ*cpaF* (electroporated plasmid pNPTSAbΔ*cpaF* into strain YB434) | This study |
| YB9232 (Ab Δ*cpaF* pAb*cpaF*) | *Asticcacaulis biprosthecum* C19 Δ*hfsA pilA*$^{G36C}$ Δ*cpaF* pMR10Ab*cpaF* (electroporated plasmid from strain YB9051 into strain YB9209) | This study |
| YB9233 (Ab Δ*cpaF* pCc*cpaF*) | *Asticcacaulis biprosthecum* C19 Δ*hfsA pilA*$^{G36C}$ Δ*cpaF* pMR10Cc*cpaF* (electroporated plasmid from strain YB8487 into strain YB9209) | This study |
| YB9258 (Ab Δ*cpaF* pCc*cpaF*$^{I355C}$) | *Asticcacaulis biprosthecum* C19 Δ*hfsA pilA*$^{G36C}$ Δ*cpaF* pMR10Cc*cpaF*$^{I355C}$ (electroporated plasmid from strain YB9200 into strain YB9209) | This study |
| ***Vibrio cholerae* strains** | | |
| TND0905 (*pilB*) | E7946 SmR, Δ*lacZ*::*lacIq*, $P_{tac}$-*tfoX*, Δ*luxO*::miniFRT, ΔVC1807::ZeoR, *pilA*$^{S67C}$ | This study |
| TND0987 (*pilB*$^{M391C}$) | E7946 SmR, Δ*lacZ*::*lacIq*, $P_{tac}$-*tfoX*, Δ*luxO*::SpecR, *comEA-mCherry*, ΔVC1807::CmR, *pilA*$^{S67C}$, *pilB*$^{M391C}$ | This study |
| **Plasmids** | | |
| pNPTS138 | Litmus 38 derivative, *ori*T *sac*B; Kan$^{R}$ | M.R.K Alley |

| | | |
|---|---|---|
| pMR10 | Low-copy plasmid with lac promotor; $Kan^R$ | Mohr et al. unpublished |
| pNPTSCB13*pilA2*$^{T36C}$ | pNPTS138 containing 605 bp upstream and 575 bp downstream of *pilA2*$^{T36C}$ mutation | This study |
| pNPTSCB13Δ*hfsDAB* | pNPTS138 containing 611 bp upstream and 613 bp downstream of *hfsDAB* locus; 20 residues of HfsD and 14 residues of HfsB were left in frame on either side of the deleted region | This study |
| pNPTSCB13*cpaF*$^{K288A}$ | pNPTS138 containing 500 bp upstream and 509 bp downstream of *cpaF*$^{K288A}$ mutation | This study |
| pNPTSCB13*cpaF*::*mCherry-cpaF* | pNPTS138 containing 511 bp upstream of *cpaF* start site followed by gene encoding mCherry, a linker, and then 519 bp of *cpaF* gene | This study |
| pNPTSCB13Δ*cpaF* | pNPTS138 containing 510 bp upstream and 504 bp downstream of *cpaF* deletion region; 15 residues of the N-terminal end and 18 residues of the C-terminal end of CpaF were left in frame on either side of the deleted region | This study |
| pMR10Cc*cpaF* | pMR10 with expression of *cpaF* under control of lac promotor | This study |
| pMR10Cc*cpaF*$^{I355C}$ | pMR10 with expression of *cpaF*$^{I355C}$ under control of lac promotor | This study |
| pMR10Cc*cpaF*$^{D310N}$ | pMR10 with expression of *cpaF*$^{D310N}$ under control of lac promotor | This study |
| pMR10Cc*cpaF*$^{F244LK245R}$ | pMR10 with expression of *cpaF*$^{F244LK245R}$ under control of lac promotor | This study |
| pAb*cpaF* | pMR10 with expression of *cpaF* from Asticcacaulis biprosthecum C19 under control of lac promotor | This study |
| pNPTS138Ab*pilA*$^{G36C}$ | pNPTS138 containing 605 bp upstream and 566 bp downstream of the *pilA*$^{G36C}$ mutation | This study |
| pNPTSAbΔ*hfsA* | pNPTS vector containing upstream and downstream regions with deletion of ABI_42610 *hsfA* gene; 3-4 residues of HfsA were left in frame on either side of the deleted region | P. Caccamo |
| pNPTS138AbΔ*cpaF* | pNPTS138 containing 523 bp upstream and 515 bp downstream of *cpaF* deletion region; 15 residues of the N-terminal end and 14 residues of the C-terminal end of CpaF were left in frame on either side of the deleted region | This study |
| pET28 | Expression vector | |
| pET28CB13*cpaF* | pET28 vector containing $His_6$-CB13*cpaF* under control of IPTG-inducible promotor | This study |
| pET28CB13*cpaF*$^{I355C}$ | pET28 vector containing $His_6$-CB13*cpaF*$^{I355C}$ under control of IPTG-inducible promotor | This study |

| | | |
|---|---|---|
| pET28CB13*cpaF*$^{F244LK245R}$ | pET28 vector containing *His$_6$*-CB13*cpaF*$^{F244LK245R}$ under control of IPTG-inducible promotor | This study |
| pET28CB13*cpaF*$^{D310N}$ | pET28 vector containing *His$_6$*-CB13*cpaF*$^{D310N}$ under control of IPTG-inducible promotor | This study |

| Primer | Sequence (bold = 5' tail/restriction site or overlap for SOE PCR and HiFi assembly, <u>underlined</u> = point mutation built into primer) | Description |
|---|---|---|
| CB13pilA2T36CF1 | **ttctggatccacgat**gatgtctcgcaccgggcccgtga | Construction of pNPTS138CB13*pilA2*$^{T36C}$ |
| CB13pilA2T36CR1 | <u>ggtgccgagggTGCagacagcggtcac</u> | Construction of pNPTS138CB13*pilA2*$^{T36C}$ |
| CB13pilA2T36CF2 | gtgaccgctgtct<u>GCA</u>ccctcggcacc | Construction of pNPTS138CB13*pilA2*$^{T36C}$ |
| CB13pilA2T36CR2 | **agcttcctgcaggat**ggggccgcacccgtgaagcaat | Construction of pNPTS138CB13*pilA2*$^{T36C}$ |
| CB13ΔhfsDABF1 | **gcgaattctggatccacgat**tggaaaagctgcggaagacgctcg | Construction of pNPTS138CB13Δ*hfsDAB* |
| CB13ΔhfsDABR1 | **ggaggctgtagcgagac**cagtacttccgtgacctgctgccg | Construction of pNPTS138CB13Δ*hfsDAB* |
| CB13ΔhfsDABF2 | **aggtcacggaagtactg**gtctcgctacagcctccggcgttc | Construction of pNPTS138CB13Δ*hfsDAB* |
| CB13ΔhfsDABR2 | **aaagcttcctgcaggat**agcagacaggccccgaaggcc | Construction of pNPTS138CB13Δ*hfsDAB* |
| CB13ΔcpaFF1 | **ttctggatccacgat**acccagaagatccgcggcgcg | Construction of pNPTS138CB13Δ*cpaF* |
| CB13ΔcpaFR1 | **gccgtaatagcgggc**ggccttgggatcgccggaggc | Construction of pNPTS138CB13Δ*cpaF* |
| CB13ΔcpaFF2 | **ggcgatcccaaggcc**gcccgctattacggcctcgagcg | Construction of pNPTS138CB13Δ*cpaF* |
| CB13ΔcpaFR2 | **agcttcctgcaggat**gtgggtccggcccttggccag | Construction of pNPTS138CB13Δ*cpaF* |
| CB13NmChyCpaFF1 | **ttctggatccacgat**tgacgacgactacgagttcggcgc | Construction of pNPTS138CB13*cpaF*::*mCherry-cpaF* |
| CB13NmChyCpaFR1 | **tcgcccttgctcaccat**catctacttcttcttgaacaggcccgagaacat | Construction of pNPTS138CB13*cpaF*::*mCherry-cpaF* |
| CB13NmChyCpaFF2 | **caagaagaagtagatg**atggtgagcaagggcgaggagga | Construction of pNPTS138CB13*cpaF*::*mCherry-cpaF* |
| CB13NmChyCpaFR2 | **gagccggcggccgagccggccgagcc**cttgtacagctcgtccatgc | Construction of pNPTS138CB13*cpaF*::*mCherry-cpaF* |

| | | |
|---|---|---|
| CB13NmChyCpaFF3 | **gctcggccgccggctcgggcgagttc**atgttcggcaagcgcgacacgtcag | Construction of pNPTS138CB13*cpaF*::*mCherry-cpaF* |
| CB13NmChyCpaFR3 | **agcttcctgcaggat**gaagacccggtgcgcgccgt | Construction of pNPTS138CB13*cpaF*::*mCherry-cpaF* |
| pMR10CB13cpaFF | **taataa/gagctc**cgcaaaagacctcgatgttctcgggcct | Construction of pMR10Cc*cpaF* plasmids |
| pMR10CB13cpaFR | **taataa/gaattc**ctactccgccgcgtccagagct | Construction of pMR10Cc*cpaF* plasmids |
| cpaFI355CR1 | gcggacttcgccgacGCAgatccgttcgggacg | Construction of Cc*cpaF*$^{I355C}$ plasmids |
| cpaFI355CF2 | cgtcccgaacggatcTGCgtcggcgaagtccgc | Construction of Cc*cpaF*$^{I355C}$ plasmids |
| cpaFF244LK245RCR1 | ggtcagcttgtccttcCTTaacttccggatggtcag | Construction of Cc*cpaF*$^{F244LK245R355C}$ plasmids |
| cpaFF244LK245RCF2 | ctgaccatccggaagttAAGgaaggacaagctgacc | Construction of Cc*cpaF*$^{F244LK245R355C}$ plasmids |
| cpaFD310NR1 | gcagttcggcggcGTTttcgcaggtcacgac | Construction of Cc*cpaF*$^{D310N}$ plasmids |
| cpaFD310NF2 | gtcgtgacctgcgaaAACgccgccgaactgc | Construction of Cc*cpaF*$^{D310N}$ plasmids |
| pMR10AbcpaFF | **taataa/aagctt**tcctgagcaacctcttcaaaaagtgagcc | Construction of pMR10Ab*cpaF* |
| pMR10AbcpaFR | **taataa/gaattc**ctattccgaagcgtccagcgcttca | Construction of pMR10Ab*cpaF* |
| AbpilAG36CF1 | **ttctggatccacgat**cagcgtatggttcttttccaccgtcac | Construction of pNPTS138Ab*pilA*$^{G36C}$ |
| AbpilAG36CR1 | gtcagcgagcccgacagggtGCAgaggatcgagatgagcgcga | Construction of pNPTS138Ab*pilA*$^{G36C}$ |
| AbpilAG36CF2 | tcgcgctcatctcgatcctcTGCaccctgtcgggctcgctgac | Construction of pNPTS138Ab*pilA*$^{G36C}$ |
| AbpilAG36CR2 | **agcttcctgcaggat**agcagaagggcggccggaaa | Construction of pNPTS138Ab*pilA*$^{G36C}$ |
| AbΔcpaFF1 | **ttctggatccacgat**atccgttcggccgcgccgtttatc | Construction of pNPTS138AbΔ*cpaF* |
| AbΔcpaFR1 | **ctcgcgctcgagtcc**gcctgcgggggctgctccccca | Construction of pNPTS138AbΔ*cpaF* |
| AbΔcpaFF2 | **gcagcccccgcaggc**ggactcgagcgcgagctcgctgaag | Construction of pNPTS138AbΔ*cpaF* |
| AbΔcpaFR2 | **agcttcctgcaggat**atcatgcccaggacccaccgcgg | Construction of pNPTS138AbΔ*cpaF* |
| pET28CB13cpaFF | **taataa/catatg**ttcggcaagcgcgacacg | Construction of pET28CB13*cpaF* plasmids |

| | | |
|---|---|---|
| pET28CB13cpaFR | **taataa/gaattc**ctactccgccgcgtccagagct | Construction of pET28CB13*cpaF* plasmids |
| DOG0408 pilBM391F1 | acgcctttcttctctgggatttc | Construction of TND0987 |
| BBC2284 pilBM391R1 | agcgctacgctcgtttctgcgccaagacccggatgtggtgTGTgtcggcgaaatccgc | Construction of TND0987 |
| BBC2285 pilBM391F2 | gcagaaacgagcgtagcgc | Construction of TND0987 |
| DOG0411 pilBM391R2 | atataacgaggtggttacgcc | Construction of TND0987 |

**Supplemental data legends.**

**Table S2.** Sheet 1) List of the accession numbers identified as TadABC-like within a local database of 2554 bacterial genomes. Sheet 2) List of the accession numbers used to generate the phylogenetic tree in Figure 1.

**Table S3.** Transposon sequencing data depicted in Fig. S3.

**Supplemental Movie 1**. *C. crescentus* CB13 with labeled pili exhibiting dynamic cycles of extension and retraction.

**Supplemental Movie 2**. *C. crescentus* CB13 with labeled pili exhibiting localized mCherry-CpaF at the base of extending and retracting pili.

**Supplemental Movie 3**. *C. crescentus* CB13 with labeled pili exhibiting delocalization of mCherry-CpaF from the base of retracting pilus that coincides with cessation of retraction.

**Supplemental Movie 4**. *C. crescentus* CB13 expressing *cpaF*$^{I355C}$ with labeled pili exhibiting slowed pilus extension and retraction.

**Supplemental Movie 5**. *C. crescentus* CB13 expressing *cpaF*$^{F244LK245R}$ with labeled pili exhibiting slowed pilus extension and retraction.

**Supplemental Movie 6**. *C. crescentus* CB13 expressing *cpaF*$^{D310N}$ with labeled pili exhibiting slowed pilus extension and retraction.

**Supplemental Movie 7**. *A. biprosthecum* expressing its own *cpaF* (AbcpaF) harboring non-dynamic, labeled pili.

**Supplemental Movie 8**. *C. crescentus* CB13 expressing *A. biprosthecum cpaF* (AbcpaF) harboring mostly non-dynamic, labeled pili.

**Supplemental Movie 9**. *A. biprosthecum* expressing *C. crescentus cpaF* (CccpaF) with labeled pili exhibiting dynamic cycles of extension and retraction.

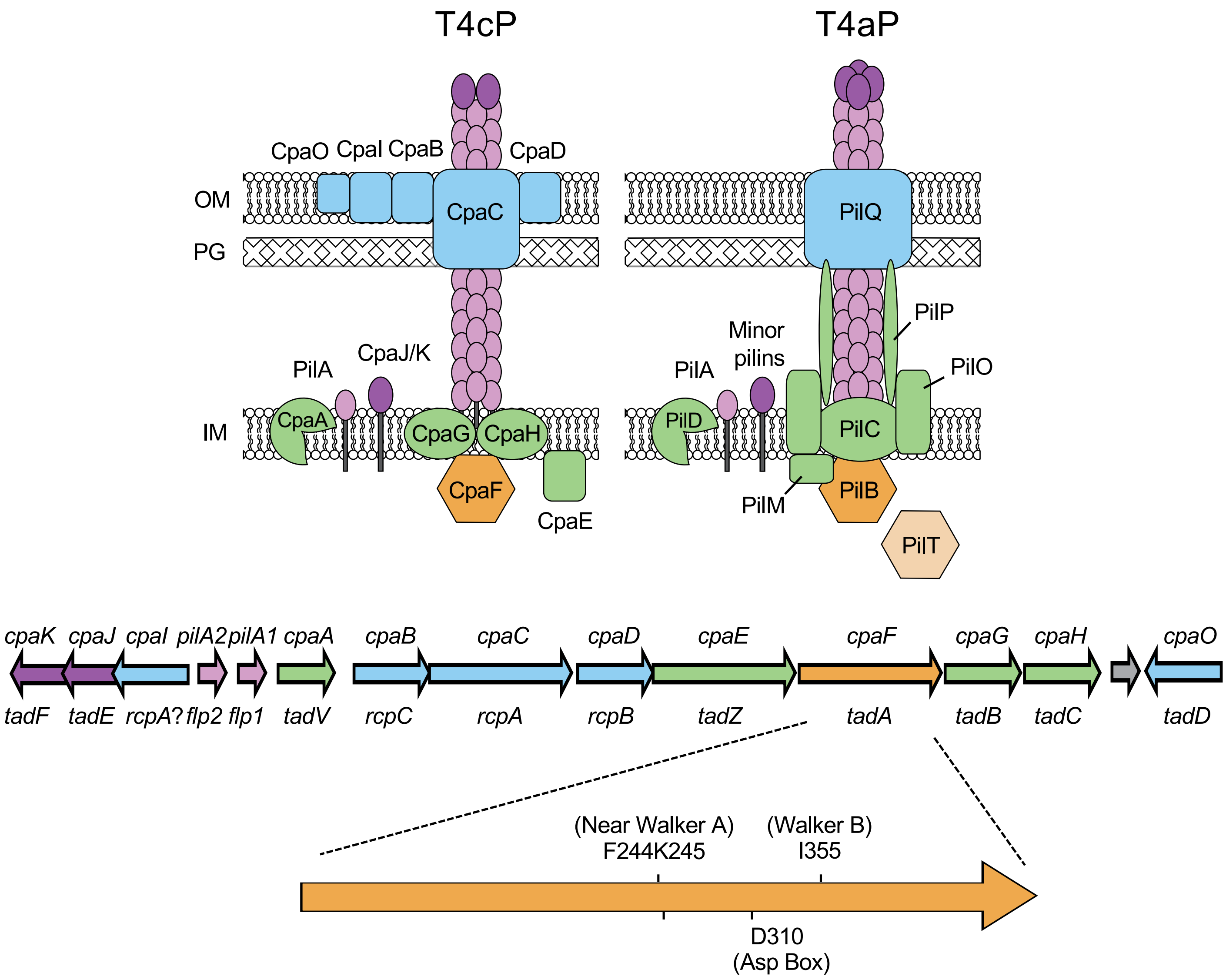


Figure S1. The tad pilus structure and gene locus in *Caulobacter crescentus* CB13. (Top) Predicted machinery structure of the tad pili (left) and type IVa pili (right). OM = outer membrane, PG = peptidoglycan, IM = inner membrane. (Bottom) Gene locus encoding components of the predicted tad pilus machinery in *Caulobacter crescentus* CB13. Top gene names are from *Caulobacter* nomenclature, bottom gene names are from original tad pilus nomenclature and ref (Mignolet et al 2018). The major pilin subunits, *pilA*, are shown in pink. A cysteine knock-in mutation in the *pilA2* subunit revealed labelable pili as seen in Figure 2 and can be sterically obstructed by the addition of PEG-mal. The orange arrow is a zoom in on the *cpaF* gene with areas of importance labeled.

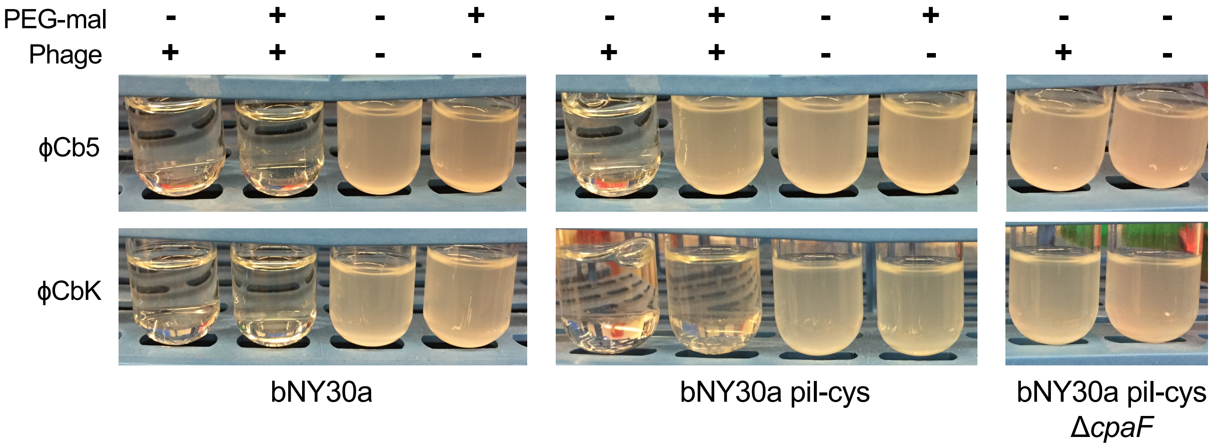


Figure S2. ϕCb5 phage require pili and their retraction for *Caulobacter* infection. Phage sensitivity assays performed in tubes on indicated strains (bNY30a pil-cys = YB9034; bNY30a pil-cys Δ*cpaF* = YB8482) with or without added 500 µM of PEG-mal to block pilus retraction or a high MOI of ϕCb5 phage (top) or ϕCbK phage (bottom). Turbid tubes indicate cell growth and phage resistance. Blocking retraction prior to phage treatment confers more resistance to ϕCb5 than ϕCbK.

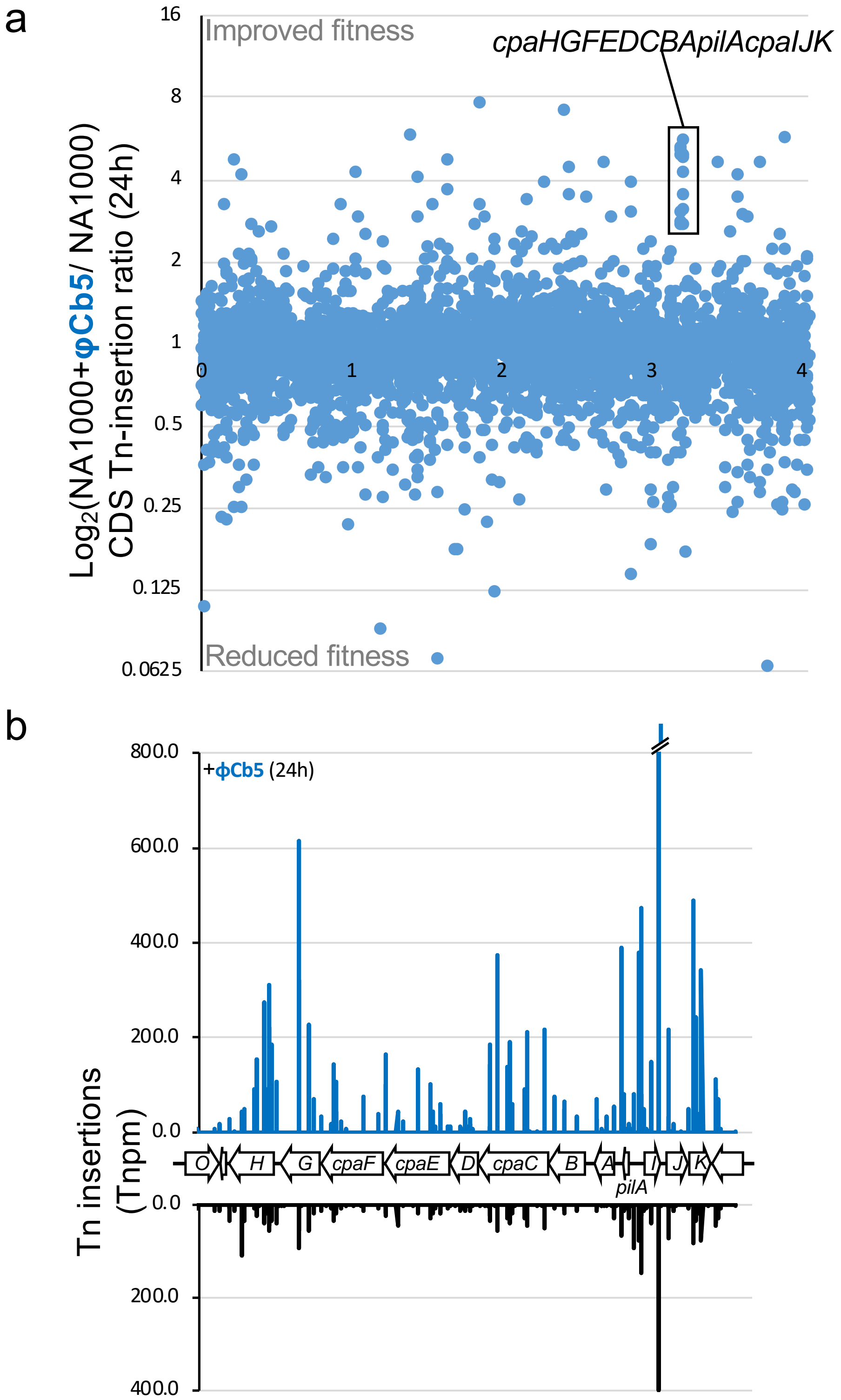


Figure S3. Tn-seq experiments reveal that Tn insertions in the pilus operon improved growth fitness during ϕCb5 phage infection in *C. crescentus* NA1000. (a) Whole genome coverage of changes in Tn-insertion ratios. (b) Tn-insertion bias at the *cpaABCDEFGHpilAcpaIJK* locus of the NA1000::Tn(Gent$^R$) bank after 24h of growth in PYE medium in absence (black) or in presence of φCb5 phage at MOI 10^3 (blue). Tn-insertion per million value (Tnpm normalized unit) is given on the y-axis of the graph. See Supplementary Table S3 for complete list and values.

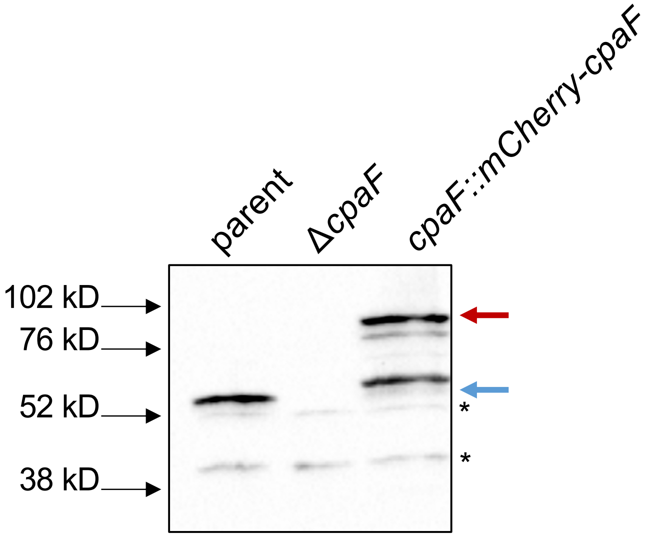


Figure S4. mCherry-CpaF is partially degraded. Western showing degradation of mCherry-CpaF protein using α-CpaF antibody. Asterisks indicate nonspecific background bands. Blue arrow indicates CpaF. Red arrow indicates expected size of mCherry-CpaF.

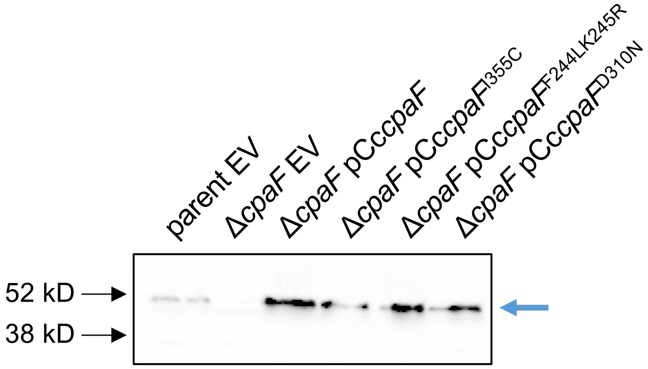


Figure S5. Mutant *cpaF* expression profiles. Western showing expression of *cpaF* in indicated strains after probing with α-CpaF antibody. Blue arrow indicates CpaF.

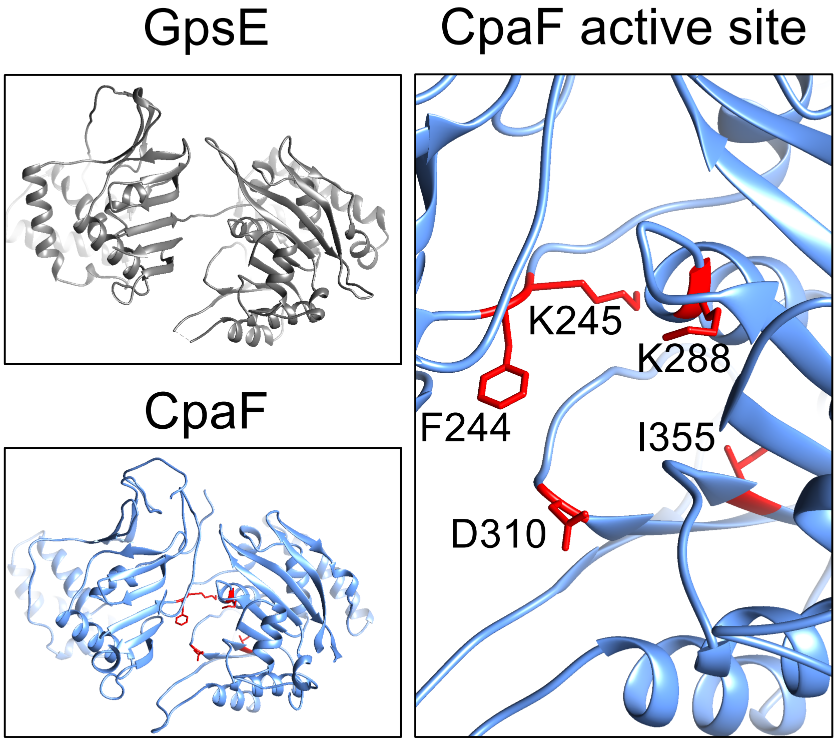


Figure S6. Mutations in *cpaF* fall into the ATPase active site of the protein. Lefthand panels are structures of ATPase monomers. 90% of *Caulobacter crescentus* CB13 CpaF (bottom left panel) was modeled at >90% confidence, revealing structural homology to archaeal type II secretion ATPase GpsE (PDB: 2OAQ, top left panel) using Phyre2 (Kelly et al. 2015) structural prediction software. Right panel shows a zoomed in perspective on the active site of CpaF with important residues including the Walker A ATP-binding lysine shown in red. Images were generated using Chimera (Petterson et al. 2004).

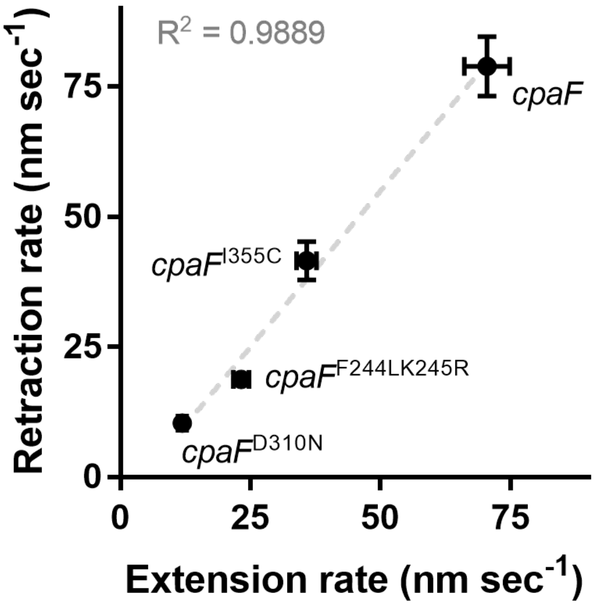


Figure S7. Extension and retraction rates of *cpaF* mutants are correlated. Correlated averages of extension and retraction rates for each mutant from (Fig. 3b). Error bars show SEM.

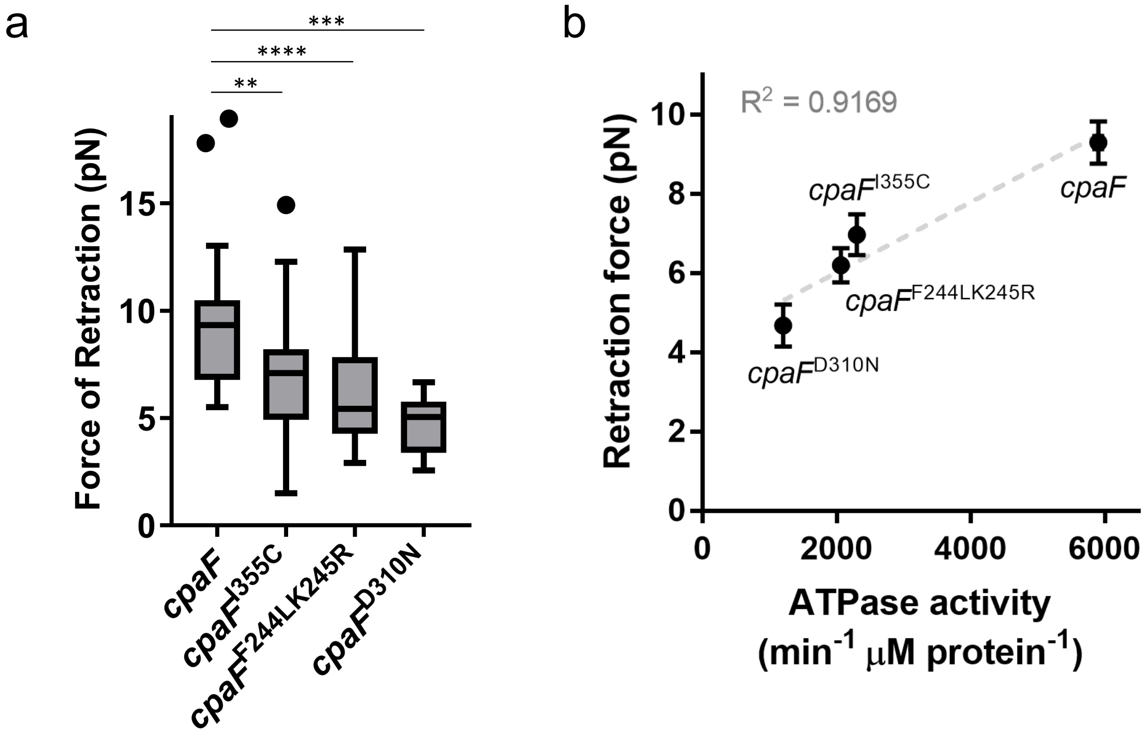


Figure S8. Forces of retraction are reduced and correlated with ATPase activity of *cpaF* mutants. (a) Retraction force measurements of indicated strains determined from micropillar assays. Box and whisker plots show Tukey's confidence intervals. *cpaF* n = 33, *cpaF*$^{I355C}$ n = 34, *cpaF*$^{F244LK245R}$ n = 34, *cpaF*$^{D310N}$ n = 7. Statistics were determined using Sidak's multiple comparisons test. P**<0.005, P***<0.001, P****<0.0001. (b) Correlated averages of retraction forces and ATPase activity for each mutant from Fig. 3. Error bars show SEM.

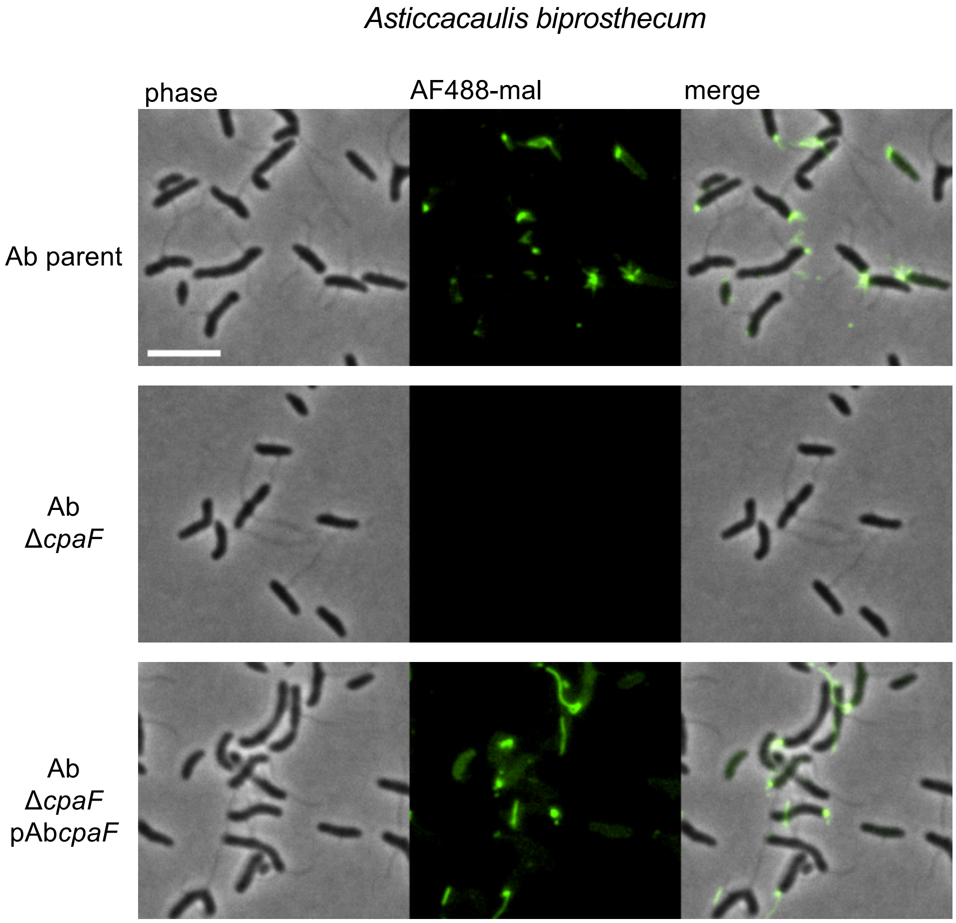


Figure S9. *Asticcacaulis biprosthecum* CpaF is required for pilus synthesis. Representative images of indicated cysteine knock-in pil-cys strains that have been labeled with AF488-mal. Scale bar is 5 μm. Unlike *C. crescentus* which harbors a single, polar stalk, *Asticcacaulis biprosthecum* harbors two stalks on the lateral sides of the cell and are visible as filaments under phase contrast.
